# Techniques and challenges in studying photosynthetic diversity in freshwater aquatic vascular plants

**DOI:** 10.1101/2025.09.24.677520

**Authors:** Ellie Loew-Mendelson, David Wickell, Calista Mrozinski, Elizabeth Doan, Jason Hupp, Richard L. Vath, Lingling Yuan, Karolina Heyduk

## Abstract

While the photosynthetic diversity of aquatic plants rivals that of terrestrial species, the environmental conditions underlying that diversity fundamentally differ. Despite these environmental differences, aquatic and terrestrial plants have convergently evolved Carbon Concentrating Mechanisms (CCMs). However, characterization of these pathways in submerged plants has lagged behind terrestrial systems due to methodological constraints of the aquatic environment. Here we review and evaluate contemporary methods for detecting CCMs in aquatic plants. Physiological methods including gas exchange and isotopic analyses provide valuable insights in terrestrial plants but face significant challenges in aquatic systems. Biochemical assays of organic acid accumulation reliably detect CCMs in aquatic species but may struggle to detect weak or non-canonical CCMs in submerged plants. Finally, we propose a method for targeted molecular assays of phosphoenolpyruvate carboxylase (PEPC) expression that could provide a sensitive tool for characterizing photosynthetic diversity in a wide range of aquatic species. Our results show that methods and frameworks developed for terrestrial plants do not necessarily directly translate to aquatic systems. However, by extending these methods and integrating multiple lines of evidence we can improve our ability to characterize photosynthetic diversity in aquatic plants.

## INTRODUCTION

Photosynthesis is central to life on Earth, converting sunlight and carbon dioxide into organic molecules that provide energy to nearly every living organism on the planet. The most common photosynthetic pathway in green plants—C_3_ photosynthesis—is characterized by Rubisco fixing carbon from CO_2_ directly into a 3-carbon compound that is eventually converted to sugars to sustain plant growth. The carboxylation efficiency of Rubisco is reduced in low carbon environments—for example, when stomata close in response to drought, CO_2_ in the leaf becomes limited. The reduced availability of CO_2_ drives an increase in a competing reaction where free oxygen is fixed in place of carbon, leading to an energetically costly process called photorespiration. Plants living in environments that promote photorespiration have frequently evolved mechanisms to minimize it (Hagemann et al., 2016; Sage and Stata, 2021). Since the initial description of C_3_ photosynthesis by Calvin, Benson and Bassham in 1948 (Calvin and Benson, 1948) and subsequent discovery of oxygenation by Rubisco (Bowes et al., 1971), characterizing the diverse ways that plants minimize photorespiration has become a central goal of modern plant physiology.

Among the processes that minimize photorespiration, carbon concentrating mechanisms (CCMs) are of interest for their ability to saturate Rubisco with CO_2_ despite environments that limit carbon. The two most common CCMs, Crassulacean Acid Metabolism (CAM) and C_4_, are derived forms of photosynthesis remarkable for their convergent evolution in every major lineage of vascular plants (Sage et al., 2011; Gilman et al., 2023). Both CAM and C_4_ use phosphoenolpyruvate carboxylase (PEPC) to fix atmospheric CO_2_ into 4-carbon intermediates, typically malate, that serve as carbon reserves. PEPC activity is separated temporally by CAM plants, where CO_2_ fixation by PEPC occurs at night and stored malate is decarboxylated during the daytime. In the C_4_ pathway, PEPC activity is separated from Rubisco spatially by sequestering PEPC in the mesophyll cells and Rubisco in specialized bundle sheath cells or, in the case of a few single-celled C_4_ plants, within intracellular compartments (Voznesenskaya et al., 2001, 2002). In both C_4_ and CAM, CO_2_ is subsequently released from the 4-carbon intermediate and concentrated around Rubisco, thereby allowing Rubisco to be consistently saturated with carbon, even when CO_2_ supply is otherwise environmentally limited.

CAM and C_4_ are complex traits involving multiple pathways that seem to arise via the co-option and regulatory rewiring of genes in existing pathways (Reyna-Llorens and Hibberd, 2017; Heyduk et al., 2019; Wai et al., 2019). In part due to this complexity, neither can be understood as a single binary trait but rather as a dynamic suite of traits that can vary in degree, timing, and context (Engelmann et al., 2003; Lauterbach et al., 2017; Winter, 2019). Trait variation is particularly true of CAM plants, which span a continuum from weak and facultative forms to strong constitutive CAM. This complexity means that the presence or absence of CAM cannot always be inferred from a single assay. Additionally, due to its environmental plasticity, plants may need to be examined under different environmental conditions to distinguish their photosynthetic pathway. Detecting and classifying CCMs therefore requires multiple lines of evidence, potentially integrating physiological, biochemical, and genetic data.

CAM and C_4_ were initially characterized in terrestrial plants (Thomas, 1949; Kortschak et al., 1965; Winter and Smith, 1996; Sage, 2004), and model plants in CCM research are overwhelmingly terrestrial species (Sage et al., 2011; Gilman et al., 2023), with the exception of the non-vascular algal CCMs (see: Beardall and Raven, 2020). Subsequent research has shown that both CAM and C_4_ are present in vascular aquatic species (Madsen and Sand-Jensen, 1991; Keeley, 1998; Bowes et al., 2002). This notable case of evolutionary convergence is evidence of a common selective pressure in both terrestrial and aquatic environments: reduced availability of CO_2_ to Rubisco. In terrestrial systems, CO_2_ limitation is typically due to water limitation or high temperatures. Low water availability can reduce stomatal conductance and limit the supply of CO_2_ inside the leaf, and high temperatures alter the kinetics of Rubisco such that it begins to favor oxygen over CO_2_ thereby increasing photorespiratory rates. In aquatic environments, poor diffusion of CO_2_ through the water column, its rapid depletion during the day by algae, and the predominance of bicarbonate over dissolved CO_2_ present similar stresses on Rubisco in submerged plants.

Terrestrial plants have independently transitioned to aquatic habitats an estimated 97 times (Meseguer et al., 2022), and many of these lineages exhibit novel photosynthetic strategies that do not map neatly onto terrestrial definitions of CAM or C_4_. For instance, terrestrial CAM species are typically defined by a tight association between stomatal regulation to reduce water loss and accompanying succulence; aquatic CAM plants do not generally exhibit either of these traits. Likewise, terrestrial C_4_ is defined by specialized Kranz anatomy to facilitate the separation of PEPC activity, but submerged aquatic C_4_ species typically lack Kranz anatomy, and typically maintain C_3_ and C_4_ pathways within the same cell (Bowes, 2011), a variation that is only occasionally seen in terrestrial plants (von Caemmerer et al., 2014).

Differences between aquatic and terrestrial adaptations often mean that assumptions and best practices underlying physiological measurements of terrestrial plants cannot be directly applied to aquatic systems, posing a unique challenge for researchers studying CCMs across diverse ecological contexts or for those interested in aquatic plant physiology. In combination with a high degree of plasticity exhibited by many aquatic plants (described in full below), it is likely that CCMs are being misclassified, underestimated, or entirely overlooked in aquatic plants using conventional approaches. With a fuller understanding of how aquatic plants assimilate carbon, we can begin to answer questions about the convergent evolution of this trait, ancestral state of photosynthesis, and the effects of photosynthetic traits on community ecology for aquatic plants. For example, did the shift of photosynthetic pathways happen before or after the transition to the aquatic habit? And, are alternative carbon assimilation pathways associated with other traits, e.g., invasivity, or resilience to environmental perturbations?

In this review, we examine the physiological, biochemical, and molecular methods that have been applied to identify CCMs in submerged aquatic vascular plants, highlighting their strengths and limitations. We assess whether methods employed for terrestrial organisms can provide meaningful information for aquatics, review a rich history of aquatic plant ecophysiology, and provide new data on molecular methods that can bridge the current gaps in studying and characterizing aquatic CCMs. We further highlight the importance of evaluating CCMs in submerged aquatic plants without being confined to the framework of past investigations in terrestrial systems. By providing a comprehensive overview of considerations and methods for investigating non-model aquatic plant physiology, we aim to facilitate continued research on the diversity of aquatic plant photosynthesis.

### Aquatic characteristics that impact physiology

Vascular aquatic plants rely on the aquatic habitat in order to complete their lifecycle, living partially or completely submerged in water. Vascular aquatic plants are distinguished taxonomically from their algal relatives, which are no less fascinating in their biology or their CCMs, but are not discussed here (for a review of CCMs in algae, see: Giordano et al., 2005; Beardall and Raven, 2020). Vascular aquatic plants represent 1.5% of angiosperm species and span 407 genera, yet arose through a remarkable 97 independent transitions from terrestrial to aquatic habits (Meseguer et al., 2022). Adaptations to the challenges of submerged life have resulted in enormous variation in aquatic plant form. Of particular interest are variations within leaf form and function, as leaves are the main site of photosynthesis. Leaves for aquatic species can range from emergent, floating, to submerged. Within submerged leaves, leaf morphology can be whole, highly dissected, or even fenestrated. The internal structure of leaves can be a few cells thick, or have large internal air spaces called aerenchyma. Submerged aquatic plants have leaf arrangements that can be defined by two categories: [1] caulescent, with long leafy stems (e.g., *Elodea, Egeria* (Hydrocharitaceae)) and [2] rosette, with ribbon-like or needle-like leaves (e.g., *Vallisneria* (Hydrocharitaceae)*, Littorella* (Plantaginaceae)*, Isoetes* (Iseotaceae))(Sculthorpe, 1967)(Fig. 1A-G).

**Figure 1.**
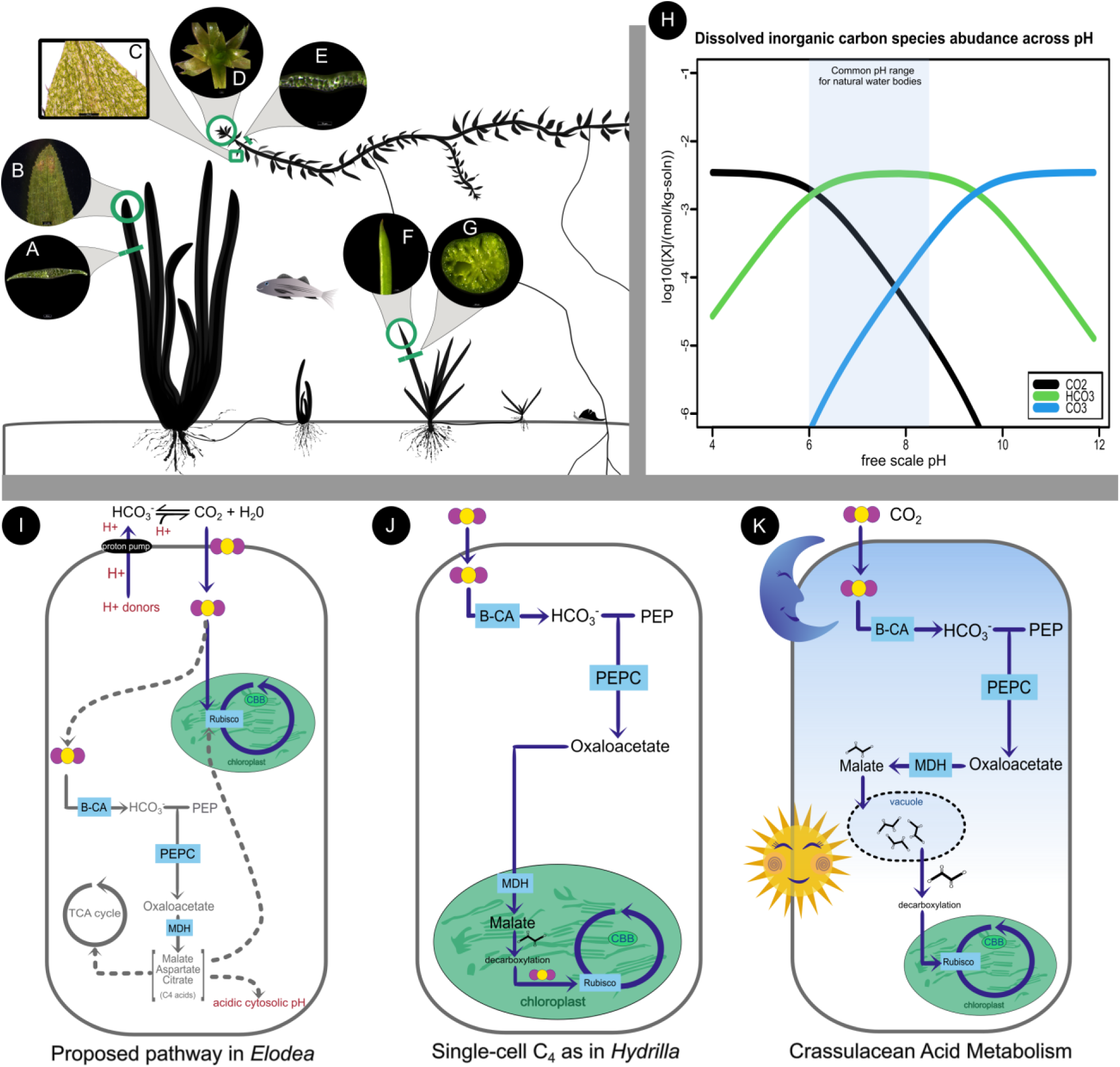
A-G) Examples of submerged aquatic vegetation forms. A) Cross section of *Vallisneria sp.* B) Adaxial leaf of *Vallisneria sp*. C) Adaxial leaf of *Egeria densa.* D) Rosette growing tip of *Egeria densa.* E) Cross section of *Egeria densa* leaf. F) Leaf surface of *Littorella uniflora.* G) Cross section of *Littorella uniflora.* H) Bjerrum plot showing the speciation of acid-base systems as a function of pH created using AquaEnv R-package (Hofmann et al., 2010). I-K) Simplified metabolic pathways for possible CCMs in aquatic species. I) Proposed pathway for *Elodea canadensis* as described in (Degroote and Kennedy, 1977). Proton pump for apoplastic conversion of HCO ^−^ to CO is depicted with H+ in red. Gray dashed lines are potential pathways for C_4_ acids. J) Single-cell C_4_ as exemplified by *Hydrilla verticillata* (modified from (Bowes et al., 2002), where C_4_ is chloroplastic. K) Crassulacean Acid Metabolism, where the sun and moon illustrate carbon uptake at night and decarboxylation during the day.

Leaves of submerged aquatic plants often have a lower density of stomata relative to terrestrial plants, or may lack functional stomata entirely (Sculthorpe and Sculthorpe, 1967; Iida et al., 2016; Horiguchi et al., 2019; Li et al., 2019). Gas exchange without stomata occurs passively, with dissolved CO_2_ diffusing directly across cell membranes (Maberly and Spence, 1983). Without functional stomata, aquatic plants cannot actively regulate gas exchange and thus do not experience the same trade-off as terrestrial plants do with regards to carbon acquisition and water-loss (Bowes et al., 2002). Uptake of CO_2_ from the atmosphere can be determined by concentration gradients rather than time of day. For example, in environments where CO_2_ is more abundant at night (e.g., heavily vegetated lake environments with nocturnal respiration) passive diffusion may result in uptake of carbon at night, which is then stored in C_4_ acids for later decarboxylation and refixation (Van et al., 1976; Holaday and Bowes, 1980).

Aquatic plants are also adapted to low carbon environments, as CO_2_ diffuses 10^4^ times more slowly in water versus air and its abundance fluctuates both seasonally and daily (Stumm and Morgan, 2013). Dissolved inorganic carbon (DIC) originates from both minerals (e.g., sediments) and CO_2_ from the atmosphere. In the water, DIC exists as multiple molecular species that are buffered in equilibrium with one another; the fluctuation between carbon dioxide (CO_2_), bicarbonate (HCO_3_-) and carbonate (CO_3_^2^-) is described as the carbonate system (Fig. 1H). A plant’s access to carbon is mediated by equilibria both within the water and between the water and the atmosphere. In slow moving or unstirred water (e.g., river water), for example, CO_2_ abundance can be very high compared to atmospheric CO_2_ (Stumm and Morgan, 2013). The pH of waterbodies largely determines the relative makeup of dissolved inorganic carbon species (although temperature and salts contribute as well) (Fig 1H). Many waterbodies where plants can be found exist at pH values where HCO_3_^−^ predominates (Fig 1H). However, CO_2_ is the only form of dissolved carbon that can passively diffuse across plant membranes, meaning that HCO_3_^−^ is not a readily accessible form of carbon for plants. Direct bicarbonate uptake is not a passive process in plants, which led to scientific debates in the early 1900’s about how aquatic plants might facilitate movement of HCO_3_-across membranes (Wilmott, 1921; James, 1928). It is now widely agreed that submerged vascular plants can acquire carbon via micro-acidification of the surrounding layer around the leaf, converting HCO_3_^−^ to CO_2_ so that it can freely diffuse across membranes (also called “apoplastic conversion”) (Cavalli et al., 4/2012; Smith and Walker, 1980; Prins et al., 1982; Rubio et al., 2017; Poschenrieder et al., 2018; Li et al., 2025). In select marine seagrasses, active transport of HCO_3_^−^ may occur through an H+ energized symport, a membrane protein (Rubio et al., 2017). Algae and cyanobacteria also have well documented HCO_3_^−^ transport mechanisms, but no such transport has been documented in freshwater vascular plants (Walker et al., 1980; Giordano et al., 2005; Beardall and Raven 2020).

The aquatic habit is clearly different from a terrestrial plant’s environment, resulting in vast differences in how submerged plants access and use CO_2_. Understanding these processes in aquatic plants and how they vary in accordance to the environment is important for describing aquatic ecosystem dynamics and, more broadly, documenting the planet’s biodiversity. Testing submerged aquatics’ photosynthetic physiology requires methods that take into account the differences in aquatic versus terrestrial environments.

### Isotope discrimination and carbon uptake

Carbon naturally occurs as two stable isotopes, ^12^C and ^13^C. These isotopes occur at different abundances in the air, with the ^13^C isotopologue of CO_2_ comprising only a small fraction of atmospheric CO_2_ (∼1%). Carboxylating enzymes discriminate between ^12^C and ^13^C to varying degrees, resulting in quantifiable differences in stable isotope ratios (δ^13^C) of ^13^C/^12^C relative to a carbon standard (Farquhar et al., 1989). Rubisco discriminates against ^13^C, while PEPC does not, resulting in a less negative δ^13^C value (i.e., closer to atmospheric) for plants that use a greater proportion of PEPC fixation (Osmond et al., 11/1973). For typical C_3_ plants, this value ranges from −22‰ to −32‰, while CAM and C_4_ are typically greater than −20‰. While carbon isotope ratios do not differentiate between C_4_ and CAM (both pathways use PEPC), this method remains a powerful tool because fresh or dry leaves can be used as the source material, including samples from herbaria (see: (Crayn et al., 2015) for a study of more than 1800 species). Due to the ease of obtaining material and the relative accessibility of stable isotope labs, carbon isotope ratios have been used widely in the terrestrial plant literature to document the diversity of C_4_ and CAM across plant lineages (Silvera et al., 2010; Horn et al., 2014; Heyduk et al., 2016)

While carbon isotope ratios are a straightforward method for delineating species that use CCMs in terrestrial plants, they are not a viable method for aquatics. Unlike terrestrial plants, aquatic plants exhibit a broader and more variable range of isotope values from −11 to −50‰ in freshwater ecosystems (Keeley and Sandquist, 1992). The broad range of ratios is in part because the source of carbon in aquatic systems is different than in the atmosphere. Carbon can either become available from diffusion via air, breakdown of minerals, or via respiration from aquatic biota, generating a bias for which isotopes may be present in water bodies. Additionally, carbon isotopes are incorporated at different rates during the conversion between carbon species, meaning that δ^13^C may contain signals of carbon source as opposed to reflecting discrimination by carboxylating enzymes (Mook et al., 1974). One area where signal of carbon source might be useful is delineating between plants that uptake bicarbonate and those that do not; for example, a study by (Maberly et al., 1992) shows a relationship between bicarbonate uptake and δ^13^C ratios where plants able to take up bicarbonate tended towards less negative values (−11.03 to −21.40‰) whereas primarily CO_2_ users had very negative δ^13^C values (−30.0 to - 34.5‰). Likewise, (Chappuis et al., 2017) sampled plants across different water bodies and found that submerged aquatic plants ranged in their δ^13^C from −43.1 to −7.5‰. They note that the variability was highest in the pH range between 7 and 8, when bicarbonate is the most common carbon form. While there might be interest in using δ^13^C to understand carbon dynamics (e.g., using CO_2_ versus HCO_3_-), this metric cannot be used to distinguish between Rubisco and PEPC carboxylation in aquatic species.

### Gas exchange and carbon assimilation rates

Direct measurement of CO_2_ uptake is routinely used in terrestrial plant physiology, and the ability to track CO_2_ uptake in the dark has been key to describing CAM species and their ecophysiological dynamics (Nobel, 1976; Black and Osmond, 2003). Such methods are typically not destructive and can be done *in situ*, allowing measurements on plants under a variety of natural and experimental conditions. For submerged aquatic species, there is an added complication that measurements must be conducted while the plant is submerged. And while we can measure the ambient CO_2_ concentration in air, direct measurements of dissolved CO_2_ in water are difficult, due to pH dependent interconversion with HCO_3_^−^, low rates of gaseous diffusion, and steep environmental gradients of CO_2_ near the boundary layer of leaves. Historically, researchers have solved these issues in measuring CO_2_ uptake in aquatic plants by bubbling air with a known concentration of CO_2_ through a chamber containing a plant sample. Infrared gas analyzers (IRGAs) were then used to measure the change in CO_2_ concentration in air of the chamber over time (Van et al., 1976; Holaday and Bowes, 1980; Keeley and Bowes, 1982). Others in the field measured oxygen evolution as a proxy for photosynthetic rates, most commonly via Clark-type O_2_ electrodes (Van et al., 1976; Colmer and Pedersen, 2008). Oxygen evolution, however, can only be a proxy of Rubisco-driven carboxylation and thus does not inform on the presence or activity of a CCMs.

Measuring gas exchange has been largely standardized for terrestrial plants thanks to the development of commercial gas analyzers, many of which are ‘open’ systems where a continuous stream of air is moved across the leaf sample (e.g., LI-COR LI-6800, Lincoln, NE USA). In such systems, the difference in CO_2_ and water vapor between the air stream before and after it interacts with a leaf are used to compute photosynthesis and transpiration rates (Busch et al., 2024). The amount of CO_2_ in the air stream is measured by an IRGA, and water vapor can be measured either via IRGAs or relative humidity sensors. These systems are powerful tools to measure terrestrial carbon assimilation, and modifications to chamber design have allowed measurements on entire plants (Huber et al., 2018), soils (Macias et al., 2025), or even ecosystems (Baldocchi, 2020). Gas analyzers have become a routine tool in terrestrial plant ecophysiology, but the nature of aquatic plants requires modifications to the chamber to accommodate a water column. LI-COR’s aquatic chamber (6800-18, LI-COR Environmental, Nebraska, USA), for example, improves upon earlier work that used a flask to submerge an aquatic plant. In the case of the LI-COR chamber, however, air is constantly circulated across the water column in an open system and the gas differentials in the headspace are used to estimate assimilation rates. The advantage of such an open gas exchange system is that it allows researchers to obtain measurements on equilibrated samples, even if it takes some amount of time for the plant sample and media to come to equilibrium. The built in fluorometer also provides a mechanism to measure photosystem function at the same time as gas exchange.

Net assimilation takes into account both photosynthesis and respiration. The CO_2_ compensation point (Γ, units: uL CO_2_ /L, L = liter) is the CO_2_ concentration at which the rate of respiration matches the rate of photosynthesis, making the net assimilation zero. The CO_2_ compensation point can help differentiate between C_3_ and C_4_ plants, the latter of which have lower Γ due to PEPC’s carbon concentrating activity (Krenzer et al., 1975). The environment heavily influences Γ in aquatic plants. To illustrate this, (Salvucci and Bowes, 1981) incubated submerged plants in two different environments to mimic summer and winter (heat/high light; cold/low light) which resulted in considerably lower Γ during the heat/high light treatment. *Hydrilla verticillata* has a Γ of 10 uL/L during the heat/high light treatment whereas during the cold/low light treatment, the Γ is 84 uL/L. *Egeria densa* showed a similar pattern with 17uL/L versus 43 uL/L in the summer vs. winter treatments. Aquatic plants have the ability to shift Γ in response to environment regardless of PEPC activity; for example, *Cabomba caroliana* is reported to have Γ values of 10 uL/L during heat/high light and 82 uL/L during cold/low light, but PEPC activity is consistently low across treatments, suggesting that submerged macrophytes may be able to modulate Γ through other means (Salvucci and Bowes, 1981). No recent work has investigated what might be driving plasticity in Γ from freshwater submerged plants, but we are hopeful that with new advances in IRGA technology further work can be done to uncover more of the unique mechanisms that aquatic plants use to prevent respiration and adapt to their low carbon environments.

#### Method testing: Gas exchange

The LI-COR aquatic chamber (6800-18, LI-COR, Nebraska, USA) has successfully been used to measure CO_2_ assimilation rates in green algae (Hupp et al., 2021; Yoshida et al., 2023), but no data has been peer-reviewed on its use for submerged freshwater aquatic vascular plants (but see unpublished thesis by (Anderson, 2024) on *Myriophyllum* and (Davey and Lawson, 2024) for a marine macrophyte). Here we tested the ability of the LI-COR 6800-18 to detect carbon flux in *Elodea canadensis* and *Egeria densa,* two submerged vascular plants. We first tested our chamber on two species of *Tetradesmus* algae (CCAP 276135 and *T. obliquus*). The LI-COR 6800-18 was fitted with a sample adapter kit (LI-COR 9968-338) that allowed for larger samples to be inserted in the side, rather than through the small opening at the top. For algal samples, we used Bold’s Basal Medium bubbled with ambient air for at least 30 minutes prior to use in the LI-COR. Pre-bubbling allowed the medium to equilibrate more quickly to chamber conditions, which approximated ambient air CO_2_ concentrations. We first verified the blank medium had zero CO_2_ flux after a 10 minute equilibration period. We then measured light response curves in replicate for both algal species, using a minimum wait time of 5 minutes and a maximum of 10 minutes before logging a measurement, and with a 30 s averaging time. We did not add carbonic anhydrase to our media (though see others who have, e.g., Hupp et al., 2021). CO_2__s was set to maintain 420 μmol mol^−1^, H_2_O control regulated H_2_O_s at 20 mmol mol^−^¹, and we used a flow rate of 500 µmol s^−^¹. Neither temperature of the solution nor pH were controlled in our trial experiments. Finally, we used 500,000-1 million cells of algae in the chamber, as estimated via a hemocytometer, in 12 mL of media and were able to detect positive flux that increased with increasing light intensity (Fig. 2AB) (measured using the light response curve program on the LI-6800, using increasing Qin light levels from 0 to 2000 μmol m^−2^ s^−1^) (Appendix S1; see Supporting Information with this article).

**Figure 2.**
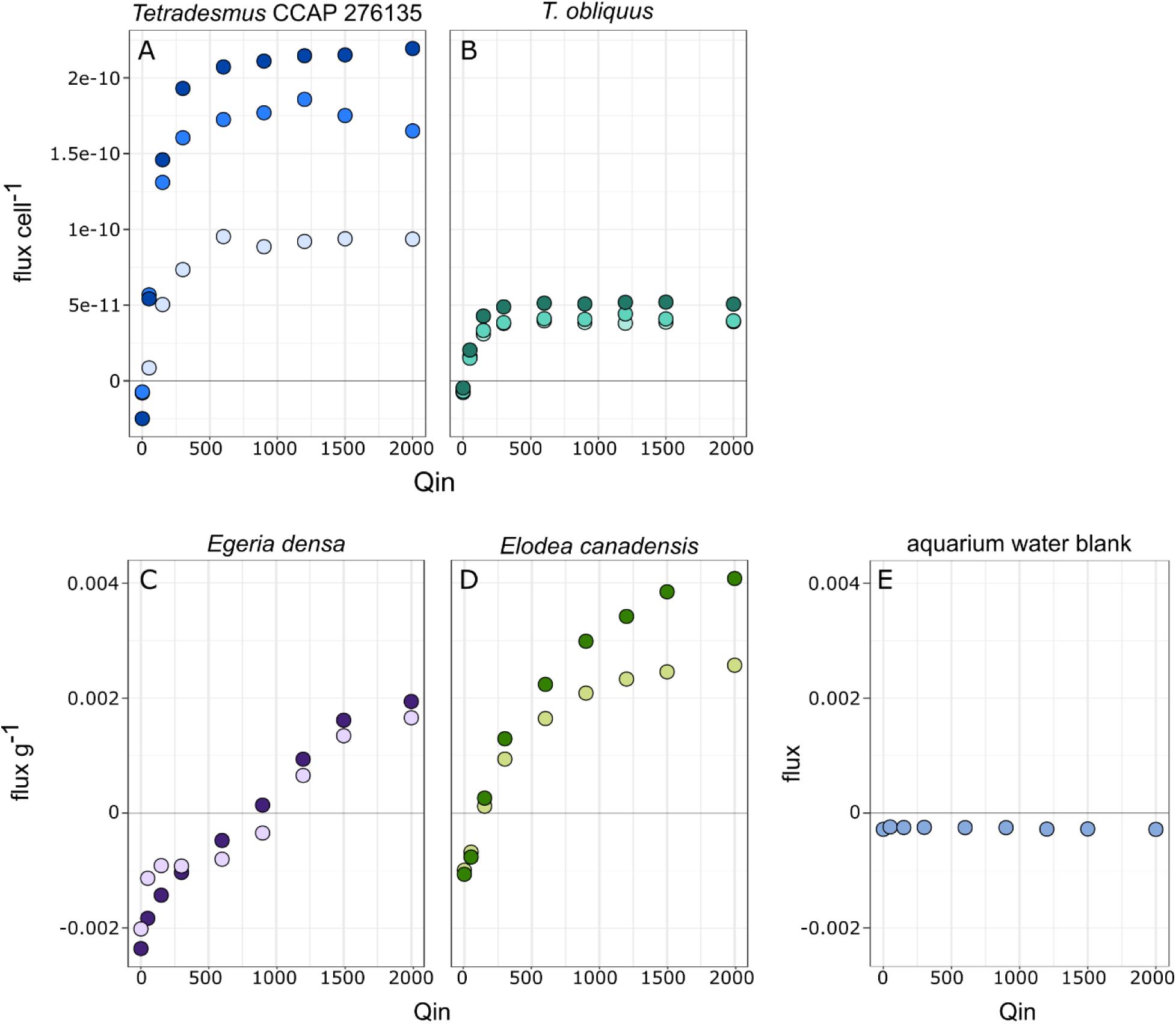
Light response curves from two species of algae (AB, top row) and two vascular submerged angiosperms (C and D, bottom row) taken at 420 μmol mol^−1^ in the Li-COR aquatic chamber. Flux is shown per cell or per mg of tissue for algae and angiosperms, respectively. Blank aquarium tank flux representing the media used for the aquatic plants is shown in panel E.

For submerged vascular aquatic plant species *Elodea canadensis* and *Egeria densa*, we targeted >1 g of tissue; amounts less than this resulted in inconsistent and negative flux values. Plants were collected from tanks with diverse species assemblages that are a part of the UConn Botanical Conservatory’s teaching collection. Our media for our aquatic vascular species was water from an aquarium containing *Litorella uniflora,* which was being bubbled continuously with ambient air and was measured as a blank before further trials (Fig. 2C). While the LI-COR aquatic chamber came equipped with rubber bands to mount the specimen on a bracket, due to the sample volume we instead packed leaf and stem material into the sample chamber without attaching it to the bracket; the density of tissue prevented excessive movement in the water column. Plants were freshly harvested before immediately being placed into the chamber. Plants that had been harvested even as recently as an hour prior to measurements no longer had positive CO_2_ fluxes, even when stored wet (e.g., in a plastic bag or beaker with water). To each plant sample in the chamber we added 12 mL of aquarium water. Plants were first starved in the chamber by providing 150 μmol mol^−1^ CO_2_ for five minutes, then allowed equilibration at 420 μmol mol^−1^ CO_2_ for 5-10 minutes. All other settings were the same as described for algae above, including no added carbonic anhydrase and no regulation of the temperature or pH, except for controlling CO_2__r at 420 μmol mol^−1^. We were able to detect positive and increasing flux in response to increasing light intensity for both species (Fig. 2CD). *Egeria*’s response to increased light intensity appeared to be slower than *Elodea*’s, which increased flux quickly at light levels about 50 μmol m^−2^ s^−1^ (Fig. 2CD). (Appendix S1). *Egeria*, on the other hand, had flux values below zero until light levels increased to nearly 1000 μmol m^−2^ s^−1^; it is possible this was a measuring error, though it was repeatable in two separate replicates run on two different days (Fig. 2C).

Due to the small volume (10-15mL) of water, the aquatic chamber exceeds ambient temperature quickly, especially at higher light intensities. While the user can control the temperature of the airstream, it often cannot cool the overall chamber without experiencing high relative humidity due to condensation. Others have used a recirculated water bath attached to the sample chamber via two ports on the chamber’s bottom side (Hupp et al., 2021; Davey and Lawson, 2024). Despite these minor issues, we could repeatedly measure light response curves in the two focal species. Further work is needed to validate the ability to measure nocturnal CO_2_ uptake in species like *Vallisneria* or *Littorella*.

### Carbon labeling

Carbon labeling is a useful tool to track the assimilation and flow of carbon through plants. Pulse-chase studies feed labeled carbon to plants (formerly labeled ^14^C, modernly ^13^C, (Pang et al., 2021)) followed by tissue sampling across a range of seconds to minutes. As the labeled carbon assimilates into the plant via initial photosynthetic carboxylation reactions, the time series sampling untangles the order in which metabolites form (Jones et al., 1983). This method was foundational in the original description of the C_3_ pathway and aided the description of C_4_ in sugarcane (the original observation: Kortschak et al., 1965) and CAM (Thomas and Ranson, 1954). In typical C_3_ plants, labeled products are overwhelmingly recovered in 3-phosphoglycerate and sugar phosphates, but in the case of CAM and C_4_ plants, the majority (>70%) of the label is recovered in 4-carbon compounds (Hatch et al., 1967; Hatch, 1971)

Multiple carbon labeling experiments using aquatic plants have recovered considerable ^14^C in C_4_ acids, generally malate and aspartate (Degroote and Kennedy, 1977; Holaday and Bowes, 1980; J. A. Browse, 1980; Reiskind and Bowes, 1991). However, none of these labeling experiments found conclusive evidence of a C_4_ pathway as evidenced by the flow of labeled carbon through C_4_ decarboxylation and subsequent fixation by Rubisco. Instead, malate stays relatively stable during the light—even as a large pool—and is steadily lost during dark respiration, suggesting that the malate pool might be mostly metabolized via the tricarboxylic acid cycle (J. A. Browse, 1980); these C_4_ acid pools could simply be used for homeostasis in the cell or for temporary storage of carbon reserves (Smith and Raven, 1976; Martinoia and Rentsch, 1994)(Fig. 1I).

Carbon labeling can extend beyond metabolite formation and also indicate carbon allocation and fixation dynamics. In *Hydrilla verticillata,* labeled carbon indicated that Rubisco was more active than PEPC, and that the species fixed primarily CO_2_ instead of HCO_3_-, reporting PEPC activity of 18 umol / mg Chl hr (umol per mg-chlorophyll-hour; samples were normalized by chlorophyll abundance and time) (Van et al., 1976). Later work in *Hydrilla* reported PEPC activity of 330 umol / mg Chl hr (Holaday and Bowes, 1980) using the same methodology. One possible explanation for these conflicting measurements is that the plants were kept in different conditions; as described earlier in this review, aquatic plants’ physiological response seems to rely heavily on the environment, and should be taken into account before experimental setup and interpreting published results.

Carbon labeling technology has advanced significantly since these studies were published; newer technologies and analytical methods, like metabolic flux analysis, can provide higher resolution of metabolic turnover and product-precursor relationships (Koley et al., 2024). An exciting development of current carbon labeling techniques is the ability to integrate the data across multiple -omic scales, and we discuss this in brief below (see “Acid accumulation and malate cycling”). Future work expanding on when and where metabolites are formed will be crucial to our understanding of how carbon flows through submerged aquatic plants (and how that changes under different environments).

### Enzymes: Localization, activity and abundance

Another approach to understand photosynthetic pathway diversity in aquatic species is to track enzymes, either in their gene expression or localization, enzyme activity and kinetics, or via screens of post-translational modifications. Enzymatic methods that assay PEPC location, activity, or kinetics are available, but these often require specialized tools or equipment. Gene expression is typically measured using whole leaf or whole plant extractions, but enzyme localization at the cell level can help clarify the dynamics of single cell-C_4_ mechanisms, as in *Hydrilla verticillata* (Reiskind et al., 1989; Fig. 1J). However, this method requires raising antibodies specific to PEPC for hybridization, followed by transmission electron microscopy to visualize sites where the antibody indicates PEPC is present within cells. Alternatively, PEPC activity can be measured through combining plant extracts with PEP substrate, but this method requires access to a mass spectrometer or other liquid scintillation technology (Van et al., 1976; Boyd et al., 2015). Finally, PEPC kinetics assays are an excellent tool for measuring PEPC affinity and carboxylation rate, both of which can be diagnostic of PEPC used for C4 photosynthesis in terrestrial plants (DiMario and Cousins, 2019). However, enzyme assays require purified proteins, either via antibody capture or through purification via *E. coli* — and neither method is necessarily straightforward to establish in a lab.

An alternative to enzymatic measurements is gene expression assays, which are relatively straightforward to set up in a lab and can be used across both model and non-model species. PEPC is present in all plants and functions to replenish citric acid cycle intermediates (O’Leary et al., 2011). However, its role as the initial carboxylation enzyme in C_4_ and CAM plants results in its transcriptional upregulation relative to C_3_ taxa (Heyduk et al., 2019). Extensive transcriptomic studies have demonstrated increased expression of PEPC and, in CAM species, a shifted peak in expression to the late afternoon and evening (Ming et al., 2015; Wai et al., 2019; Wickell et al., 2021; Heyduk et al., 2022). In *Portulaca oleracea*, a C_4_ plant that can use CAM under drought, CAM marker genes are upregulated exclusively under drought, and upon rewatering are no longer upregulated (Ferrari et al., 2020a, b). The increase in transcript expression in CCM taxa could be a useful tool for screening aquatic species, which are recalcitrant to other methods (described above).

#### Method testing: Genetic expression of the enzyme PEPC

We tested the use of a gene expression assay to screen photosynthetic diversity in aquatic plants in the Hydrocharitaceae (*Egeria densa, Elodea canadensis,* and *Vallisneria sp*.). Using OrthoFinder (Emms and Kelly, 2019), we first obtained multi-sequence alignments from available transcriptomes in the Hydrocharitaceae (Chen et al., 2022) and created family specific primers for plant-type PEPC and the reference genes ubiquitin C (UBC) and actin11. We collected and flash froze plants in liquid nitrogen every 4-hours for 24-hours (6:00, 10:00, 14:00, 18:00, 22:00, 2:00), using 3 biological replicates per species. Plants were collected from tanks with diverse assemblages of species in teaching-focused greenhouses of the UConn Botanical Conservatory; available light was ambient and, at the time of sampling, dawn occurred ∼5:00 am and dusk at ∼8:00 pm. pH was measured at time of collection and ranged across the day and night in the *Elodea* and *Egeria* tanks, but was largely consistent across time in tanks of *Vallisneria* and *Litorrella* (Appendix S2). We extracted total RNA using the Monarch RNA-Spin Kit (T2110, New England BioLabs, Ipswich, MA, USA). We filtered for quality using Qubit and Nanodrop, only keeping samples with suitable 260/280 ratios between 1.8 and 2.2 and concentration values above 10 ng/uL of RNA. With 20 ng of extracted RNA as input, we synthesized cDNA with the Protoscript-II First Strand Kit via the standard protocol (E6560, New England BioLabs, Ipswich, MA, USA). The resulting cDNA was diluted to 10 ng/uL and used for quantitative PCR with SsoAdvanced Universal SYBR Green Supermix (Cat. #1725270, Bio-Rad, Hercules, CA, USA) on a Bio-Rad qPCR thermocycler (Appendix S3).

The RT-qPCR data was analyzed by species using the *2*^−*ΔΔCt*^ method (Livak and Schmittgen, 2001). We normalized expression using the 6:00 timepoint as a control in all species but used different reference genes depending on species (UBC: *Egeria, Elodea* | ACT11: *Vallisneria, Littorella*). *Vallisneria sp.* had 3-fold higher expression of PEPC at 22:00 relative to 6:00 (*t*(3.03) = 5.25, p = 0.0066)(Fig. 3A); prior work suggested species in the genus were CAM, though results were largely inconclusive due to a lack of detectable malate accumulation (Keeley, 1998). *Egeria densa*, putatively a C_4_ species, had no significant fold-change of PEPC expression across time points relative to 6:00 (Fig. 3A). We also screened *Elodea canadensis*, commonly thought to be C_3_, but which had a significant 3-fold change in PEPC expression at 22:00 relative to 6:00 (*t*(2.03) = 5.63, p = 0.0146)(Fig. 3A). Finally, we screened a known aquatic CAM species, *Littorella uniflora*. We found significant 2-fold change in PEPC expression at 22:00 relative to 6:00 (*t*(3.05) = 3.78, p = 0.0.0157); *Litorella* appeared to have a shifted peak in PEPC expression to earlier in the afternoon, with a 3-fold change in PEPC expressions at 14:00 relative to 6:00 (*t*(3.63) = 5.12, p = 0.00446)(Fig. 3A). As an initial screen for CCMs, we argue that expression based assays are more sensitive than titrations (see below, “Acid accumulation”), can provide valuable information about the evolutionary history of CCMs in a clade, and can also help describe alternative carbon flow through PEPC that is not associated with malate (see below, “Acid accumulation and malate cycling”).

**Figure 3.**
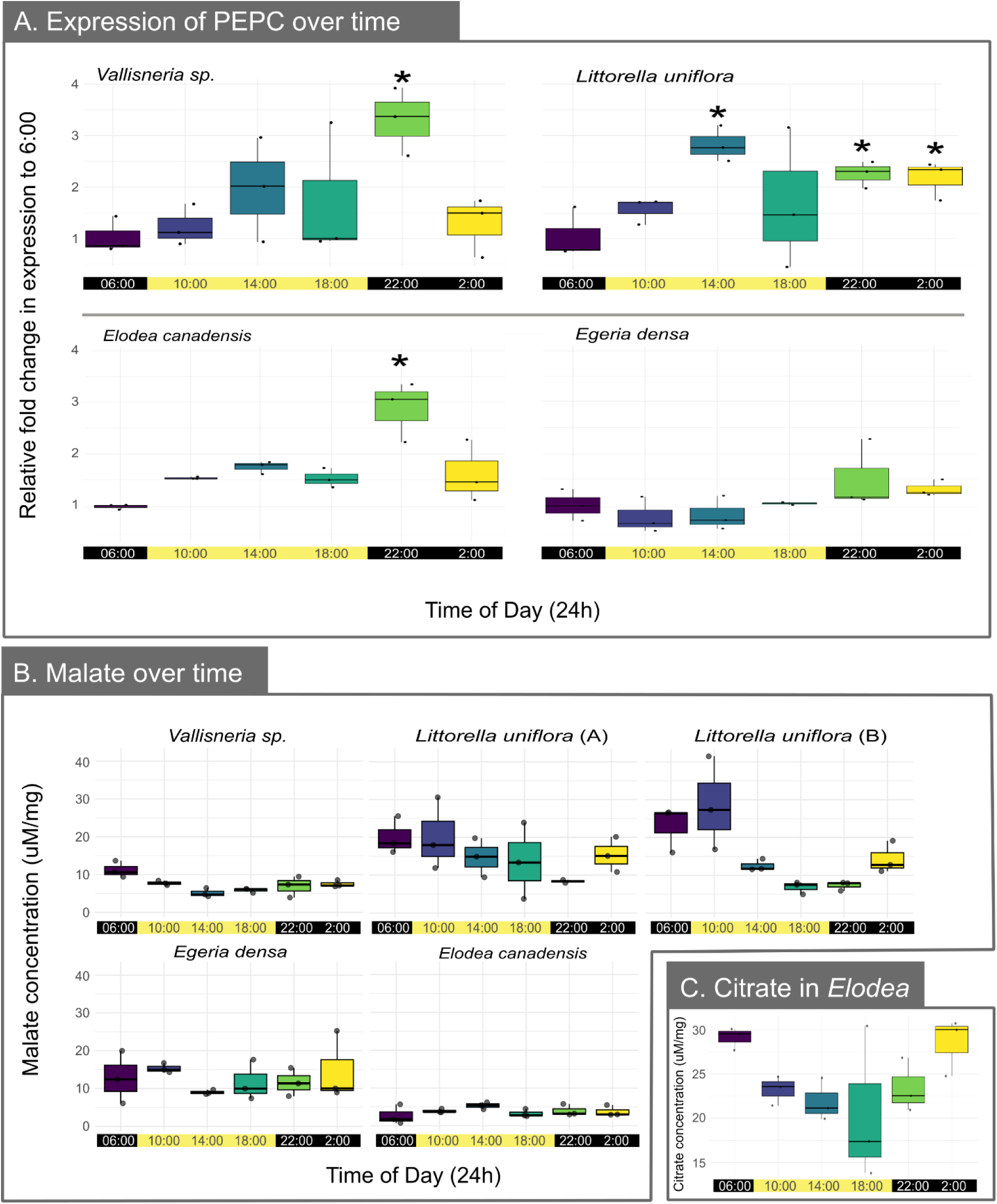
A) Expression of the enzyme PEPC over time relative to a reference gene, reported as fold-change values (mean and standard error) normalized to 6:00AM; asterisks indicate a significant fold-change at p < 0.05. B) Malate concentration over time, normalized by fresh weight. *Littorella* (A) shows results from a single leaf per collection, while *Littorella* (B) shows results from whole plants. **C**. Citrate cycling in *Elodea*.

The use of PEPC expression as a screening tool for CCMs is not without its limitations. Reference transcriptomes were available in our family of interest, allowing us to design primers with relative ease. Our focal gene, PEPC, is multicopy in most plant genomes, and as a result our RT-qPCR amplicons are likely not specific to a particular gene copy. Still, the induction in expression of the photosynthesis-related PEPC copy is likely to be strong enough to swamp the background signal of other PEPC copies that we expect to be largely constitutively expressed (Heyduk et al., 2019). We also did not target the bacterial-type PEPC copy, which has been shown to be related to CAM function in a few lineages (Heyduk et al., 2022), including in the aquatic *Isoetes* (Wickell et al., 2021). Other target genes were considered, including PEPC’s dedicated kinase, PPCK, whose activity is thought to be critical for PEPC function in a CAM and C_4_ cellular context (Taybi et al., 2000; Boxall et al., 2017). PPCK shows consistent dark-induced expression across CAM species making it a promising molecular marker for identifying CAM activity. However, PPCK is not highly conserved across taxa and we were not able to get it to consistently amplify across species within Hydrocharitaceae due to its high (∼68.7%) GC content. In some lineages, however, PPCK expression may prove a practical marker for CAM and C_4_ activity (Ferrari et al., 2020b).

### Acid accumulation and malate cycling

Overnight acid accumulation in photosynthetic tissues is a strong indicator of CAM (Fig. 1K) and can be measured through a variety of methods that range from indirect (e.g., total titratable acidity) to direct (e.g., enzymatic assays) measures of malate. Diel titratable acidity has been used to screen and flag potential CAM species across many aquatic genera (Keeley, 1998). While there are no issues with using titratable acidity as an indicator of CAM with aquatic species, there are two significant caveats. The first is that titratable acidity does not measure for malate specifically, and aquatic plants may be accumulating other acids that could mask the pattern of malate specifically (notably this is an issue for terrestrial plant measurements as well (Lüttge, 1988; Franco et al., 1992; Freschi et al., 2010). The second is that titratable acidity may not detect weak CAM phenotypes (Winter and Smith, 2022), as the acid accumulation may be very low or variable between replicates, which may be the case for many aquatic plants.

Historically, titratable acidity is used as a first pass to determining CAM activity because the protocol is relatively simple and less expensive than direct measurements of malate. However, malate is not the only acid that can accumulate in plant cells, and quantifying metabolites directly can reveal what is causing diurnal shifts in plant acidity. For example, (Keeley, 1998) reports a significant difference in morning vs. evening acid accumulation (ΔH+) for *Vallisneria americana,* but malate accumulation did not cycle diurnally (Table 1). Direct measurements of malate abundance for aquatic plants have mostly been published pre-2000, and the standard method at the time was enzymatic assays (Degroote and Kennedy, 1977; J. A. Browse, 1980; Madsen, 1987; Keeley, 1998). These assays rely on the principle that Malate Dehydrogenase catalyzes a reaction between L-malate and NAD+ to create NADH; the amount of L-malate can be calculated from the increase of NADH (Bergmeyer, 1983). This is measured via spectrophotometric absorbance but uses volatile chemicals; a safer alternative are pre-mixed colorimetric assays (e.g., MAK511 from Millipore Sigma) in which NADH reduces a colored reagent and can also be measured on the spectrophotometer.

**Table 1.** Acid accumulation measured by titratable acidity and malate assays. Averages reported with standard deviation. – : not reported Δ : change between morning and evening * : Significant according to Welch’s t-test (2-tailed, p < 0.05), ໐ : significance not reported

| **Table 1 - Acid accumulation measured by titratable acidity and malate assays**<br>Averages reported with standard deviation.<br>– : not reported $\Delta$ : change between morning and evening<br>* : Significant according to Welch's t-test (2-tailed, $p < 0.05$),<br>o : significance not reported | | | | | |
| --- | --- | --- | --- | --- | --- |
| Citation | Species | Acid<br>(mmol/kg FW)<br>18:00 | Acid<br>(mmol/kg<br>FW)<br>6:00 | Malate<br>(ug/mg<br>FW)<br>18:00 | Malate<br>(ug/mg<br>FW)<br>6:00 |
| This study - protocol A | *Egeria densa* | 0 | 0 | 12 +/- 5 | 13 +/- 7 |
| Browse 1980 | *Egeria densa* | – | – | 114 | 50 |
| Webb et al 1988 | *Egeria densa*, December | – | $\Delta 2 \pm 1$ \| o | – | – |
| Webb et al 1988 | *Egeria densa*, September | – | $\Delta 1 \pm 1$ \| o | – | – |
| This study - protocol A | *Elodea canadensis* | 0 | 0 | 3 +/- 1 | 3 +/- 3 |
| Keeley and Morton 1982 | *Elodea canadensis* | 6 +/- 3 | 5 +/- 6 | 9 +/- 1 | 16 +/- 3 |
| Webb et al 1988 | *Elodea canadensis*, May | – | $\Delta 6 \pm 4$ \| o | – | – |
| This study - protocol A | *Littorella uniflora* | 0 | 0 | 14 +/- 10 | 20 +/- 5 |
| This study - protocol B | *Littorella uniflora* | 4 +/- 1 | 49 +/- 5 \| *<sup>1</sup> | 7 +/- 2 | 23 +/- 6 \| *<sup>1</sup> |
| Keeley and Morton 1982 | *Littorella uniflora* | 19 +/- 22 | 112 +/- 14 \| * | 21 +/- 14 | 66 +/- 8 \| * |
| This study, protocol A | *Vallisneria sp.* | 0 | 0 | 6 +/- 1 | 11 +/- 2 \| *<sup>2</sup> |
| Keeley 1998 | *Vallisneria americana* | 11 +/- 7 | 42 +/- 12 * | 8 +/- 1 | 6 +/- 1 |
| Keeley 1998 | *Vallisneria spiralis* | 6 +/- 3 | 13 +/- 6 | 6 +/- 1 | 10 +/- 4 |
| Raven 1988 | *Vallisneria spiralis* | 8 +/- 1 | 9 +/- 1 | 3 +/- 1 | 3 +/- 1 |
| Webb et al 1988 | *Vallisneria spiralis*,<br>December | – | $\Delta 51 \pm 1$ \| o | – | $\Delta 54$ \| o |
| Webb et al 1988 | *Vallisneria spiralis*,<br>September | – | $\Delta 26 \pm 9$ \| o | – | $\Delta 27$ \| o |

1 - *Littorella* B: acid $p=1.052e-05$ , malate $p=0.01809$ 899 2 - *Vallisneria sp.*: malate $p=0.0218$ .

Finally, metabolomics holds great promise for understanding metabolism in aquatic plants, though perhaps not as a first pass screening tool. Metabolomics methods characterize a large swatch of metabolite abundances within organisms, but require access to a liquid chromatography or gas chromatography mass spectrophotometer (LC-MS/GC-MS). These methods, paired with labeled carbon methods, could provide insight into the first products of photosynthesis in aquatic species and thus determine whether a species is using a CCM. Additionally, the ability to track the abundance of many hundreds of metabolites may demonstrate that other metabolic pathways are important for the aquatic lifestyle, or may demonstrate entirely new pathways for carbon to flow that have not been described in terrestrial species.

#### Method testing: Titratable acidity and malate accumulation

We used previously published protocols for terrestrial plants (e.g., (Heyduk et al., 2020) and found that *Egeria densa, Elodea canadensis, Vallisneria spp.* and *Littorella uniflora* do not show diel shifts in titratable acidity (Table 1). Due to previous literature that firmly establishes *Littorella uniflora* as a plant that cycles acidity diurnally (Madsen and Sand-Jensen, 1991; Keeley, 1998), we tried a slightly different protocol with more tissue, less volume, and a homogenization step with liquid N_2_. Significant acid accumulation was detectable in *L. uniflora* with these changes (Table 1). It may be necessary to alter protocols in order to detect the low level acid accumulation in aquatic plants; for example, (Keeley, 1982; Madsen, 1987) both macerate tissue prior to titrations.

Along with titratable acidity, we performed malate assays (MAK511, Millipore Sigma, Burlington, MA, USA) on the same tissue to compare malate accumulation with titratable acidity, following kit guidelines but using half reaction volumes. Tissue for malate assays was homogenized with grinding samples in liquid N_2_ in a mortar and pestle; weight of tissue was recorded to normalize malate concentration by mg of tissue. We found that while *Vallisneria sp.* shows no difference in ΔH+ between morning and evening samples, it does cycle malate diurnally (Fig. 3B). For weak CAM species that may rely on the CAM cycle for only a small proportion of carbon gain, malate assays are particularly useful as they are much more sensitive to small diurnal changes than measuring gross titratable acidity.

*Egeria densa,* described putatively as C_3_ + C_4_ (or, C_4_-like; (Casati et al., 2000-8), had malate present without any specific pattern across time. While this could support the presence of a C_4_ cycle, where malate is produced and rapidly decarboxylated, malate is also produced by all species and used in support of the citric acid cycle. Our results from the malate assay alone cannot conclusively determine the end product of malate and thus cannot distinguish between C_4_ and C_3_.

*Elodea canadensis* did not accumulate or increase malate across the 24-hours. Following the absence of malate but a high peak of PEPC activity, we investigated citrate accumulation in *Elodea;* prior studies in CAM plants have indicated citrate cycling adjacent to the CAM cycle, but the function is unclear (Lüttge, 1990; Freschi et al., 2010). We used a citrate assay kit (MAK333, Millipore Sigma, Burlington, MA, USA) and found that citrate cycles in *E. canadensis* in a pattern similar to malate of CAM species (Fig. 3C). While other C_3_ plants do cycle citrate, their transcript abundance for PEPC does not have higher expression at night (e.g., (Scheible et al., 2000). These results highlight the importance of using multiple lines of evidence in aquatics, whose biochemistry is strongly affected by the unique aquatic environment (Appendix S3).

Overall, we found the malate kit to be more reliable and less time consuming than acid titrations for measuring acid accumulation, and it was able to detect low levels of acid cycling that were not detected via titrations (as in *Vallisneria,* Fig. 3B). However, screening the entire plant metabolome could more quickly catalogue any organic acids that may be cycling, as in the case of citrate in *Elodea*.

### Future directions

We are not the first to point out that submerged aquatic plants do not fit into the discrete categories of C_3_, C_4_ and CAM as described by terrestrial literature; in fact, most of the work reviewed here makes explicit mention of the fact that aquatic plants seem to have some but not all of the characteristics of a CCM pathway (be that C_4_ or CAM). Early biochemical work attempting to quantify photosynthetic rates in the presence of bicarbonate were also perplexed by the variation in photosynthesis depending on the environment (Wilmott, 1921; James, 1928). The ability to accurately characterize the diversity of alternative photosynthetic pathways in submerged aquatics will enable extensive research spanning biogeochemistry, ecology, evolution and physiology. Within the context of global and anthropogenic change, aquatic ecosystems have become highly nutrient dense due to chemical runoff (eutrophication). Changes in the water have led to imbalances and changes in the species assemblages, such as an increase in harmful algal blooms and expanding *Hydrilla* infestations. Aquatic plants occupy diverse roles in freshwater ecosystems, from ecologically problematic to rare. Dynamics between nuisance and native aquatic plants is a growing area of research (June-Wells et al., 2013; Lech and Willig, 2021; Miller et al., 2022) and plant physiology and photosynthesis sit at the center of processes like nutrient assimilation and plant development that mitigate dynamics between species. For example, competition between nuisance species and native species may change depending on whether they exploit the same or different carbon pools (e.g., take up carbon at night versus day). There is extensive management of weedy aquatics with herbicides; understanding of a plant’s photosynthetic physiology can help us determine more accurately how these species respond to herbicide treatment. In order to untangle how submerged plants are responding to global change, we need to establish an expectation of their current physiology. Likewise, before we can begin to untangle evolutionary origins and consequences of CCMs and their associated traits (e.g., ancestral state reconstruction, trait-trait and trait-environment correlations), we need to be able to accurately describe photosynthetic traits and how they differ among species.

Addressing gaps in methodology is the first step to describing CCMs more fully in vascular aquatic plants. The LI-6800-18 is a promising new tool to help describe photosynthetic patterns in aquatics, as the portable machine can be used in the field (with exceeding care) to obtain measurements unaffected by greenhouse tank conditions. Particularly exciting is the prospect of measuring plants in the field across an environmental gradient, or in micro-environments within a single water body and season. Higher resolution carbon labeling will also help untangle the dynamics of when and why aquatic plants form C_4_ molecules and the fate of those molecules in metabolism. Finally, we show that integrating gene expression with metabolite assays can provide a clearer understanding of flux through particular metabolic pathways; continued efforts in this area can help us better describe aquatic plant photosynthetic metabolism without the constraints of terrestrial CCM definitions.

## Acknowledgements

The authors thank the UConn Botanical Conservatory for allowing us to sample plants from their collection. This work was possible with support from the Rosalind Chair in Ecology and Evolutionary Biology endowment fund at UConn startup funding to K.H, and from the Trainor Endowment Fund and the Department of Ecology and Evolutionary Biology and Connecticut State Museum of Natural History at UConn awarded to E.L-M.

## Author Contributions

E.L-M. designed and conducted malate and gene expression assays and wrote the manuscript; D.W. assisted in data collection and contributed to writing the manuscript; C.M. conducted titratable acidity measurements; E.D. assisted with initial primer design and RT-PCRs; J.H, R.L.V, and L.Y all provided invaluable advice and guidance on the use of the Li-COR 6800-18 and contributed to writing the manuscript; K.H. guided the project, contributed to data collection, and assisted with writing the manuscript.

## Data Availability Statement

Raw LiCor, malate accumulation, and RT-PCR reaction conditions and primer sequences can be found in Supporting Information.

## Supporting Information

Additional Supporting Information may be found online in the Supporting Information section at the end of the article.

**Appendix S1** - Raw LiCor data from both algal and macrophyte measurements.

**Appendix S2** - pH of water at time of collection for malate/expression

**Appendix S3** - RT-PCR primers and reaction conditions

**Appendix S4** - Malate accumulation data

