## Appendix S2 for "Techniques and challenges in studying photosynthetic diversity in freshwater aquatic vascular plants"

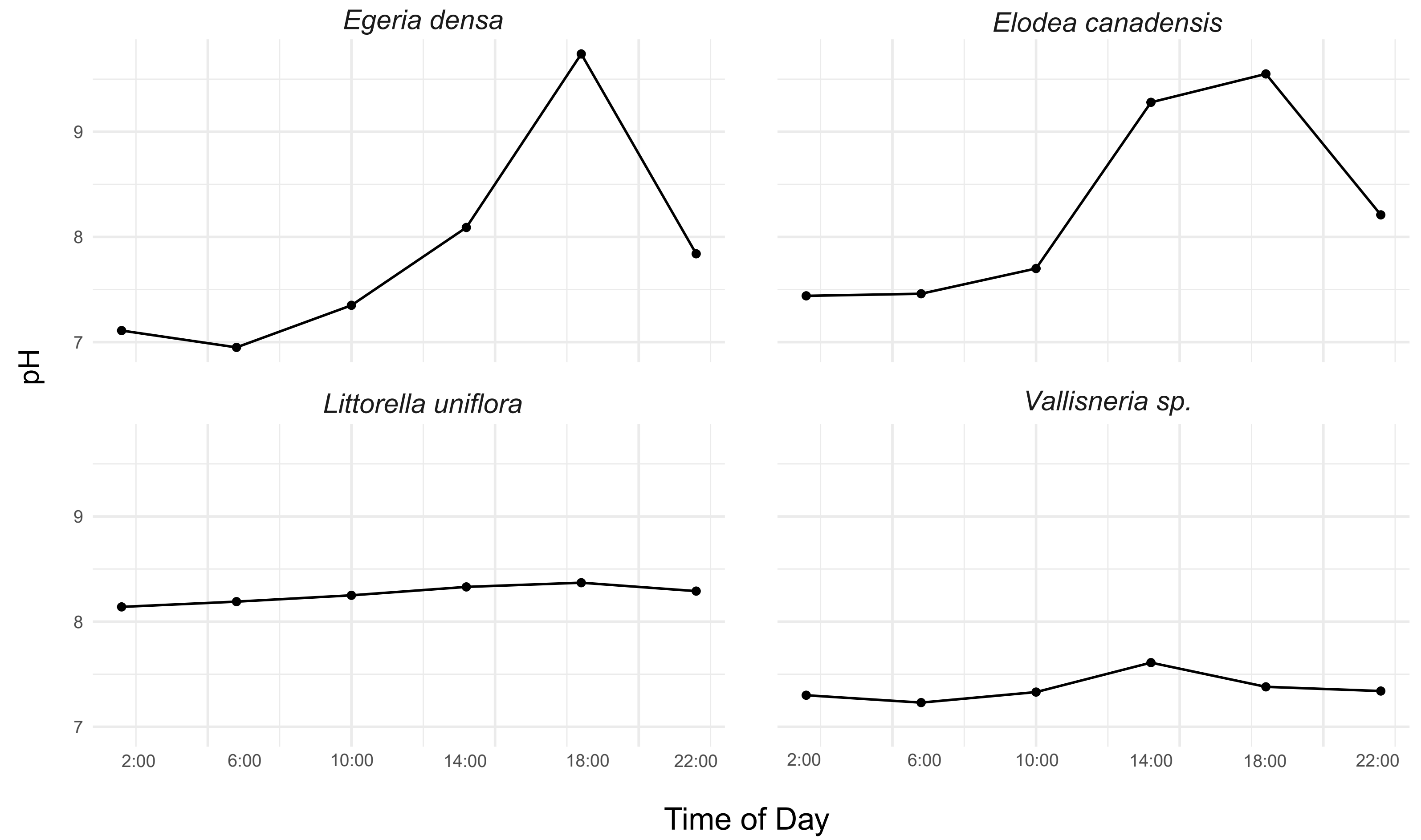


**Appendix S2** - pH of aquarium tanks where the four focal experimental species were collected, measured at each timepoint sampled for both titration/malate and qRT-PCR.
